## Supplementary file 1 for "An arms race between 5’ppp-RNA virus and its alternative recognition receptor MDA5 in RIG-I-lost teleost fish"

**Supplementary file 1.** PCR primer information in this study.

| Primer | Sequences (5'-3') |
|---|---|
| | **Primers for Real-time PCR** |
| MDA5-qRT-F | TGAGATGACTGGTGGCTTAC |
| MDA5-qRT-R | CTTCTGGCTGTTGCTACTGA |
| LGP2-qRT-F | TGAGGCGGATGTGGTGATTT |
| LGP2-qRT-R | GATGAGCAGGGCGTCGTTGT |
| STING-qRT-F | CAACGCCAACATTTCTCAC |
| STING-qRT-R | ACGCTGTGCTTGTAGACCC |
| YTHDF1-qRT-F | AGGACGTGCCAAACAGTCAG |
| YTHDF1-qRT-R | CATAGTGGGAGAAGTCATCAAAGA |
| YTHDF2-qRT-F | AGTGGCATCGACTTCTCGG |
| YTHDF2-qRT-R | CCTGGTCAAGGCTGTTCATA |
| YTHDF3-qRT-F | GGTATGAGCAGTATGGCAG |
| YTHDF3-qRT-R | GTTTGACCTTGGGTTGTG |
| METTL3-qRT-F | TGGGTAAGTTTGCTGTGGT |
| METTL3-qRT-R | GGATGGGAATGTTCAGTTTT |
| METTL14-qRT-F | GAGGAAGATGTGGAGGAAC |
| METTL14-qRT-R | AGTAGTCATTGTGCGGATT |
| IFN-1-qRT-F | TACGATGGCTAATAACTCC |
| IFN-1-qRT-R | CATTGACAAAGTGCTCCA |
| Mx1-qRT-F | GCTGCTTGTTTACTCCCA |
| Mx1-qRT-R | ACCTGCATCATCTCCCTC |
| ISG15-qRT-F | TGAACGGACAGAAGACGC |
| ISG15-qRT-R | TGAGGAATACCTGCATGG |
| Viperin-qRT-F | ACCCGTCCAAGTCCATAC |
| Viperin-qRT-R | TCATGTCAGCTTTGCTCC |
| SCRV-M-qRT-F | TCAACCTGGCAAACAACA |
| SCRV-M-qRT-R | CCTCGGACCTCTGCTTCT |
| SCRV-G-qRT-F | TCTGCCATAAGACTACCTG |
| SCRV-G-qRT-R | TCTTGACGGTGATGAATG |
| β-actin-qRT-F | GAGCCGCACGCTTCTTT |
| β-actin-qRT-R | CTGCTGTAGCCGAGGAC |
| | **Primers for MeRIP-qPCR** |
| MDA5-m6a-qRT-1F | CCCCGGCAGTAGTTCTTC |
| MDA5-m6a-qRT-1R | GACCAAAAGGAGCGGATC |
| MDA5-m6a-qRT-2F | CAAGCGGCTGTGGATCTCCT |
| MDA5-m6a-qRT-2R | ATGAAGACTGAAGATGTGCGAG |
| | **Primers for plasmid construction** |
| MDA5-exon1-GLO-F | CCGTTTAAAGGAGAAACGAAACTGAAAG |
| MDA5-exon1-GLO-R | TGCTCTAGAGGTAAGCCACCAGTCATC |
| MDA5-exon1-GFP-XhoI-F | CCGCTCGAGGGAGAAACGAAACTGAAAG |

| | |
|---|---|
| MDA5-exon1-GFP-EcoRI-R | CCGGAATTCGGTAAGCCACCAGTCATC |
| MDA5-exon1-mut1-F | GAAGTCTTTACACCGAGGCTGCGGGAGCTCGT |
| MDA5-exon1-mut1-R | CTCGGTGTAAAGACTTCAATGAGACGCACGTTTAGC |
| MDA5-exon1-mut2-F | TCGAAGCGGTCCAAAAGGAGCGGATCCTAAAA |
| MDA5-exon1-mut2-R | CTTTTGGACCGCTTCGATCAAATGTAAATAAA |
| MDA5-exon1-mut3-F | ATGCAGCGGTCTACATGCAGCGTAATATCCCG |
| MDA5-exon1-mut3-R | CATGTAGACCGCTGCATAGTGACAACCTGATT |
| MDA5-exon1-mut4-F | GAAGTCTGAAGATGTGCGAGTACACTGTCTGT |
| MDA5-exon1-mut4-R | GCACATCTTCAGACTTCATATCCATAAGACTGGGAGCC |
| MDA5-NotI-F | ATATCCATCACACTGGCGGCCGCTTCATAATGGCATCTGATAACGATG |
| MDA5-XbaI-R | TATAGAATAGGGCCCTCTAGACTGTCAACGTGATTATCAGTGTTTCTT |
| MDA5-△RD-F | AACGAGAACCCGTCTGAAGCGAGCCAGGTGGCAGAC |
| MDA5-△RD-R | TTCAGACGGGTTCTCGTTCTTCATGGTTTCTG |
| METTL3-HindIII-F | CCCAAGCTTCTCGTCATGTCGGACACAT |
| METTL3-EcoRI-R | CCGGAATTCTGCGGGGATCACATACAG |
| METTL14-HindIII-F | CCCAAGCTTGCTTCACGAAGGAGAAAAT |
| METTL14-EcoRI-R | CCGGAATTCTGTGATTGGAGATTCATAAGG |
| YTHDF1-KpnI-F | CGGGGTACCCATTTCAACATGACCACCAA |
| YTHDF1-EcoRI-R | CCGGAATTCGCAGCCGTCTTCTGTTTACT |
| YTHDF2-HindIII-F | CCCAAGCTTTGCCGCGTTTATTTAGGAG |
| YTHDF2-EcoRI-R | CCGGAATTCGGGCCATTGCAGTCTTTT |
| YTHDF3-HindIII-F | CCCAAGCTTTCAGTGCAAAACGGATCAAT |
| YTHDF3-BamHI-R | CGCGGATCCGCCTTTCCTCCTTTGTGGTT |
| FTO-KpnI-1F | CGGGGTACCCACAACTCCAGGAACATG |
| FTO-XbaI-1R | TGCTCTAGAGCCTCTACATTAAGAAAAGC |
| ALKBH5-HindIII-F | CCCAAGCTTGGCTATCTGTCAGCTACTAC |
| ALKBH5-EcoRI-R | CCGGAATTCCTTCGTCTGCCAGAAAC |
| LGP2-1F | CGGGGTACCATGTACCCATACGATGTTCCAGATTACGCTGCAGACTTTGCACTGTATG |
| LGP2-1R | GCGCCGTCTAGATCAATCAAAGAGGTCAGGGAAG |
| IRF3-KpnI-1F | CGGGGTACCATGTACCCATACGATGTTCCAGATTACGCTTCTCATTCTAAACCTCTGCTCATC |
| IRF3-XbaI-1R | TGCTCTAGAGTGTCAGTACAGCTCCATCATCTC |
| STING-HindIII-F | CCCAAGCTTCTGTGCCTCCAGGATCA |
| STING-EcoRI-R | CCGGAATTCCGATCCAGCTCGTCCTCC |
| MAVS- HindIII-F | GACGATGACGACAAGAAGCTTTCGTCTGCCAAAGACAAACTGTA |
| MAVS- EcoRI-R | TGATGGATATCTGCAGAATTCCAGCCTCTGTCCTGTCTACTTCATG |
| ggaMDA5-HindIII-F | GACGATGACGACAAGAAGCTTATGTCGGAGGAGTGCCGAG |
| ggaMDA5-XbaI-R | TGATGGATATCTGCAGAATTCTTAATCTTCATCACTTGAAGGACAATG |
| ggaMDA5-His-F | GACTACAAAGACGATGACGACAAGCATCATCATCATCATCATTAATCTAGAGGGCCCGTT |
| ggaMDA5-His-R | ATGATGATGATGATGATGCTTGTCGTCATCGTCTTTGTAGTCATCTT |

| | |
|---|---|
| | CATCACTTGAAGGAC |
| mmiMDA5-His-F | ACTTACCATCATCATCATCATCATTGAGGATCCACTAGTAACGGCCGC |
| mmiMDA5-His-R | TCCTCAATGATGATGATGATGATGGTAAGTTTCTTCCTCCTCTGAGCTG |
| | **Primers for RNA probe synthesis** |
| 5’ppp-VSV-F | TAATACGACTCACTATAGGGACGAAGACAAACAAACCATTATTATCATTAAAATTTTATTTTTTATCTGGTTTTGTGGTCTTCGTC |
| 5’ppp-VSV-R | GACGAAGACCACAAAACCAGATAAAAAATAAAATTTTAATGATAATAATGGTTTGTTTGTCTTCGTC |
| 5’ppp-SCRV-F | TAATACGACTCACTATAGGGACGAGAAAAAAGAAACCAATATACAGATTATCAATTGCTAATCAGAGACTGTGTTTGTTTTTCTCGT |
| 5’ppp-SCRV-R | ACGAGAAAAACAAACACAGTCTCTGATTAGCAATTGATAATCTGTATATTGGTTTCTTTTTTCTCGT |
| 112bp-dsRNA-F | TAATACGACTCACTATAGGGAGATGGCATCTGATAACGATGA |
| 112bp-dsRNA-R△T7 | GTAAATAAATCAGGACCTGACTCCCT |
| 112bp-dsRNA-F△T7 | AGGGAGATGGCATCTGATAACGATGA |
| 112bp-dsRNA-R | GTAAATAAATCAGGACCTGACTCCCTATAGTGAGTCGTATTA |
