## Supplementary figures and images for "An arms race between 5’ppp-RNA virus and its alternative recognition receptor MDA5 in RIG-I-lost teleost fish"

### Fig.6-figure supplement 1

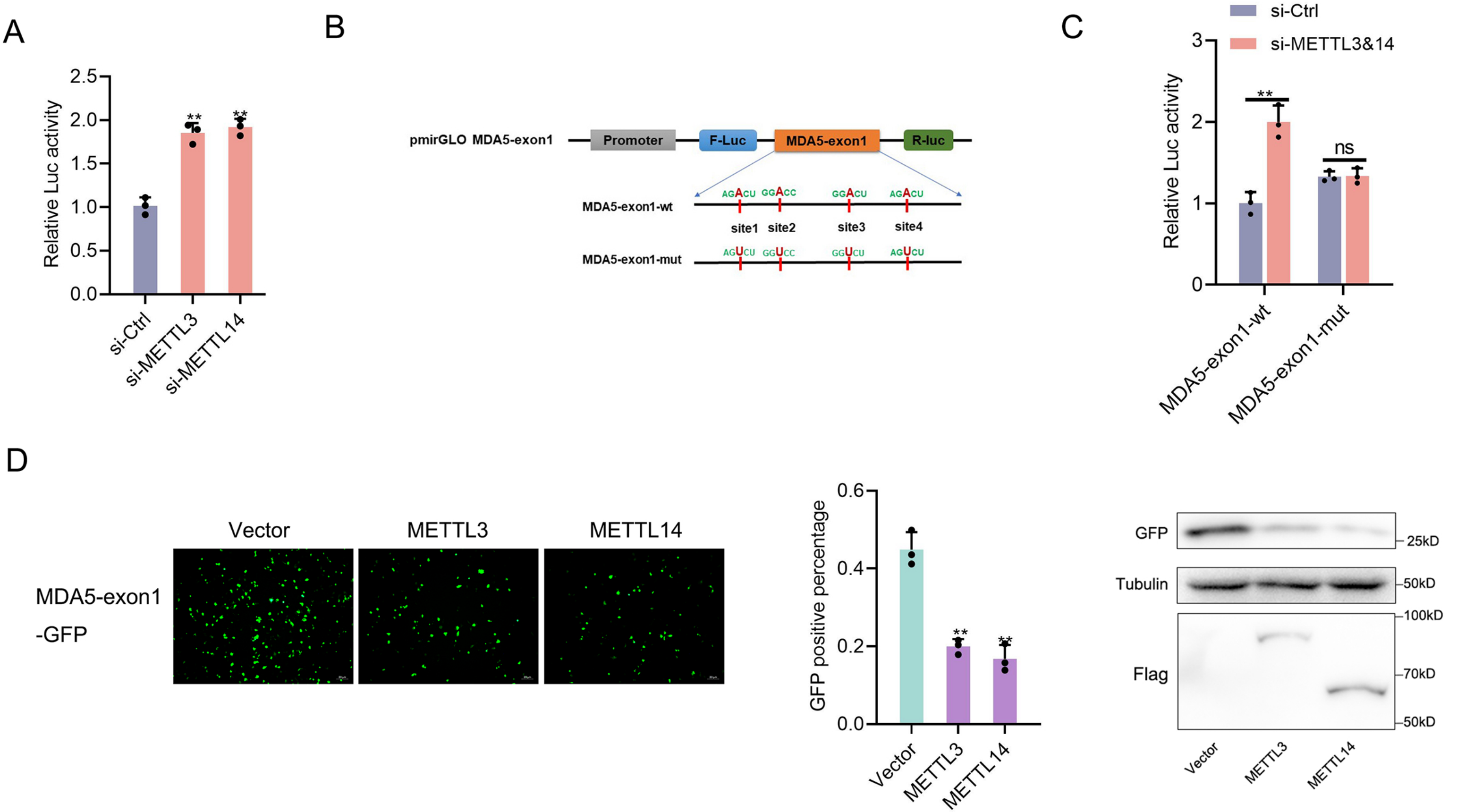

### Fig.7-figure supplement 1.

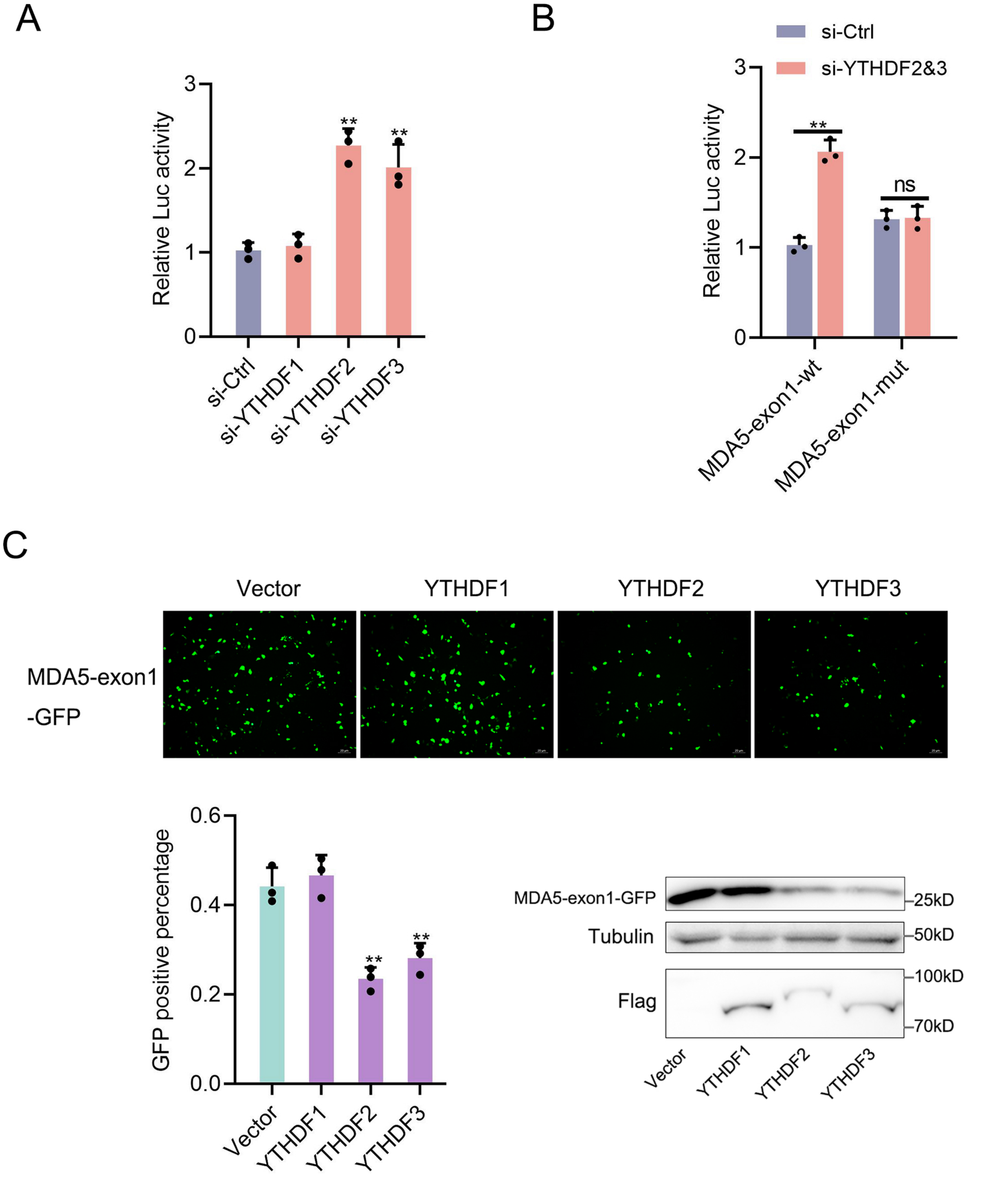
